# *In situ* investigation of extracellular vesicles in viscous formulations: interplay of nanoparticle transport and nano rheology through interferometric light microscopy analysis

**DOI:** 10.1101/2024.04.12.589108

**Authors:** Lucile Alexandre, Anastasiia Dubrova, Aruna Kunduru, Marie Berger, Imane Boucenna, Florence Gazeau, Amanda K. A. Silva, Stéphanie Mangenot, Kelly Aubertin

## Abstract

While extracellular vesicles (EVs) demonstrate growing potential as innovative therapeutics in diverse medical context (cancer, regenerative medicine, etc.) or as naturally circulant diagnostic / prognostic probes, their physical properties (size, transport, etc.) remains a critical concern. Here, we introduce a pipeline that relies on interferometric light microscopy (ILM) for measuring not only nanoparticle concentration and size distribution but also for analyzing the interactions of these nanoparticles with their environment. The analysis of interference patterns allows for the physical characterization of (bio)nanoparticles not only in aqueous solutions but also in challenging media with relatively high viscosity, particularly pertinent for characterizing gel-based EV-delivery systems. Through exploration of the instrument’s functionality and the use of calibrated NPs of various known sizes, we successfully obtained information about the local viscosity characteristics of a complex fluid embedding EVs. We present a proof-of-concept for characterizing EVs suspended in unconventional media and their interactions with their surroundings. Leveraging the outcomes of this investigation, we not only highlight the advantages of using ILM for characterizing EVs in complex fluid, particularly pertinent for the development of optimized biological carriers for targeted drug delivery and therapeutic applications, but we also validate a new method for measuring viscosity at the nanoscale.

## Introduction

Extracellular vesicles (EVs) are nanometric objects, ranging from tens to hundreds of nanometers, encapsulating proteins, lipids and nucleic acids [1]. They play a crucial role in intercellular communication, traveling between the secreting cell and the recipient cell, where they can release their cargo [2]. The transported molecules reflect the biological signature and environmental cues of the cell from which they originate. It has been shown that EVs participate to various biological phenomena, such as the embryonic development [3] or the therapy resistance of certain cancers [4], or even the initiation pro-metastatic niches distant to primary tumor [5]. For that, EVs must cross complex, sometimes media with particular mechanical properties (e.g.: high viscosity). EVs travel either by crossing the extracellular environment of tissues or by being transported by biological fluids. This transport occurs through the whole body, exhibiting a half-life of less than 30 min *in vivo* across most tissues [6]. It requires crossing porous environments, where the size of the pores (down to 20 nm) can be smaller than the average size of the vesicles [7,8] (hundreds of nm). This transport relies on the properties of both the matrix and EVs, depending on either one of them (diffusion, deformation of constrictions) or on both (molecular interactions, release of active principles). However, the transport properties of EVs are rarely studied and the dynamics governing this transport are not well-explored [8,9].

In a pathological context, the abundance, size, biomolecular composition, and biological signatures of EVs position them as inherent diagnostic probes in body fluids, serving as stable repositories of disease biomarkers [10]. Due to the inherent heterogeneity of their cargo, the isolation and analysis of EVs yield a comprehensive reservoir of diverse molecular information, affording them a comparative advantage over other biomarkers and potentially enabling the early detection of diseases. This diagnostic prowess has been demonstrated across various conditions, including colorectal [11], pancreatic [12], lung [13], and breast [14] cancers, as well as Parkinson’s [15] and Alzheimer’s [16] diseases. The potential of characterizing the physical properties of EVs in complex biofluids without any purification process was only poorly explored and could lead to potential new biomarkers discovery. In tandem with these applications, attention has turned towards EVs as nanocarriers for drugs [17], capitalizing on their inherent attributes. The lipid bilayer carrying and protecting their cargo, coupled with their targeting properties, and their transport capabilities in complex media, positions EVs as promising candidates for the effective transport and delivery of therapeutic agents.

Recent attention has been directed towards exploring EVs as promising pro-regenerative nanotherapeutic agents for tissue repair [18,19]. This interest stems from observations in successful cell therapy scenarios where administered cells do not persist, prompting a shift in understanding therapeutic effects attributed to external factors, specifically cell secretions, notably EVs. Numerous studies have demonstrated the advantageous outcomes of EV administration in the regeneration of diverse tissues such as the heart [20], kidney [21], liver [22], lung [23], brain [24], bones [25] and skin [26]. Notably, our team has focused on the local application of EVs derived from mesenchymal stem cells (MSCs) within a thermo-actuated poloxamer 407 for fistula healing, thereby facilitating the targeted delivery of EVs and contributing to the healing process [27,28]. These studies have not only substantiated the therapeutic potential of EVs for tissue regeneration but have also engendered new inquiries into the role of EVs in tissue organization, their transport through viscous matrices, and their interactions within their environment.

There are several challenges in the attempt to answer these questions, notably the absence of a standardized instrument for EV *in situ* characterization in complex media. The nanometric scale of EVs, falling below the limit of diffraction, precludes standard optical microscopic imaging technique. In response, diverse instruments have been developed to face this challenge. Transmission Electron Microscopy (TEM) and Cryo-Electron Microscopy (CryoEM) are powerful techniques for the characterization of EVs at the nanoscale[29]. However, as TEM has to be performed on dehydrated sample, the cause of its usual donut shape on images, it precludes any size and morphology measurement. With this limitation, CryoEM appears as a good alternative, as it preserves EVs in a near-native state by rapidly vitrifying the sample, providing detailed insights and is particularly valuable for studying EVs with delicate structures. However, (cryo-)TEM are expensive, not easily accessible techniques with low transferability yield into the clinics. The Nanoparticle Tracking Analysis (NTA) methodology, relying on Brownian motion analysis, enables real-time quantification of particle concentration and determination of hydrodynamic radius following particle tracking within a small observation chamber. The nano flow cytometry gives access to the same information but relies on rapidly flowing fluid stream measurement. These technologies are limited to characterizing EVs within a very low viscous matrix (typically viscosity of water). Additionally, Atomic Force Microscopy (AFM) facilitates high-resolution imaging and mechanical mapping of EV surfaces, contributing to a comprehensive understanding of their (bio)physical properties, but is limited by its throughput. However, there is still a lack of instrumental support to study the interactions of EVs within their environment.

Previously described instruments rely on the use of the Stokes-Einstein equation (Eq. 1):

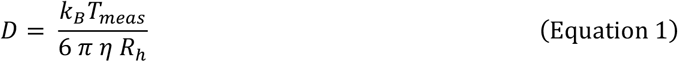

Where *D* is the diffusion coefficient, *k*_*B*_ the Boltzmann constant, *T*_*meas*_ the temperature measured during the experiment, *η* the dynamic viscosity and *R*_*h*_ the hydrodynamic radius of the particle. At a constant temperature and viscosity, the diffusion coefficient is inversely proportional to the size of the particle. Measuring the diffusion patterns of the particles in a solution enables access to the size distribution of particles within the sample, as depicted in the inset of Figure 1C. However, this assertion relies on a precise measurement of the temperature and of the viscosity. Those two measurements should be performed in the local environment of the analyzed particles. The measurement of the local viscosity at the surface of the diffusion object is a challenge in the field of soft matter, as it required access of a nano-probe. When working with hydrogels or biological matrices, viscoelastic properties can span a wide range, with Young’s modulus ranging from a kPa to several MPa. These properties exhibit a strong dependence on factors such as the matrix’s crosslinking, composition, and environmental conditions. Furthermore, the size of the object used to probe the mechanical properties in such complex medium can influence the value of the measured parameters. Atomic force microscopy (AFM) is a well-established tool for assessing the local viscoelastic properties of soft materials. Complementary, resonance force microscopy (CR-FM) methods offer a quantitative estimation of the viscoelastic moduli of polymer blends. However, both technologies require substantial instrumentation, highlighting the necessity for simpler and more accessible measurement techniques.

**Figure 1:**
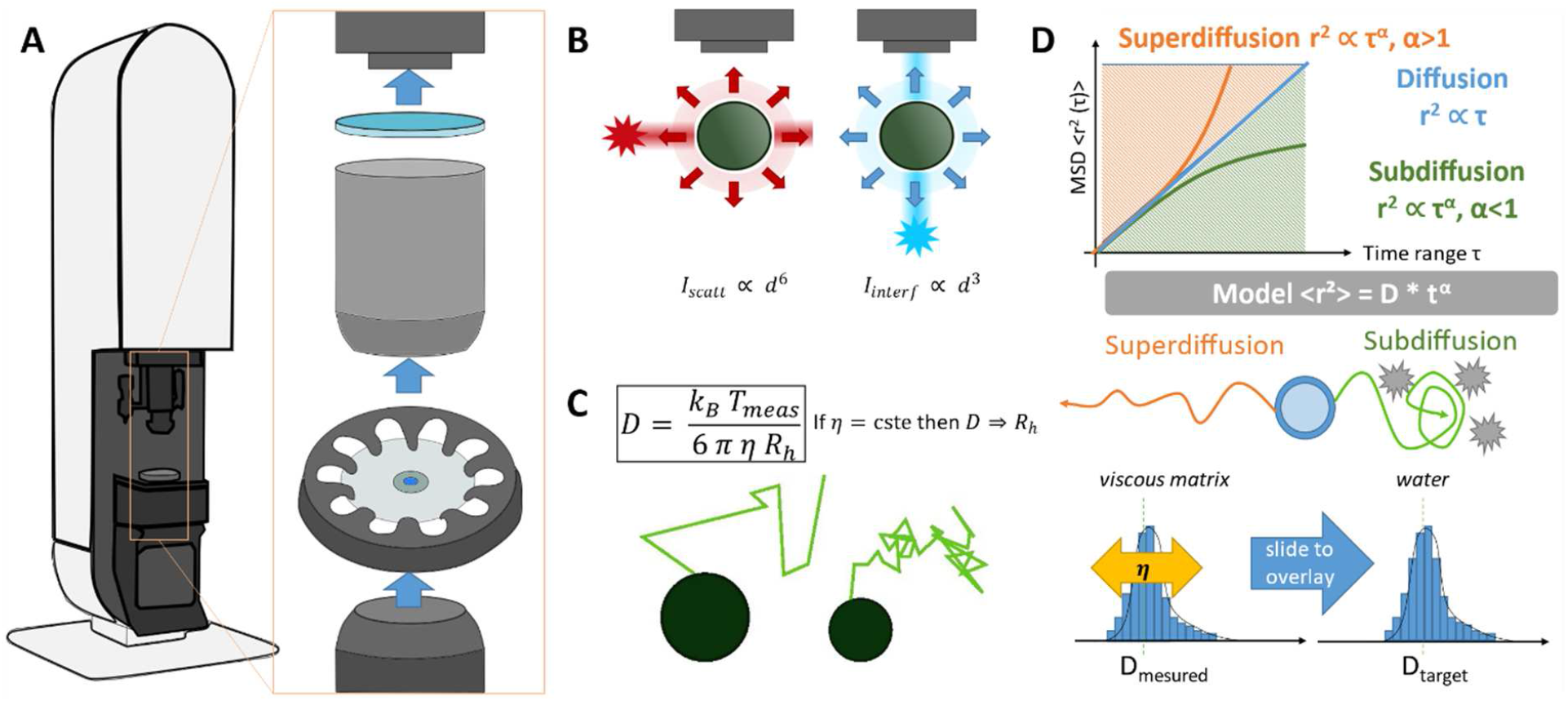
Presentation of the ILM instrument for nanoparticles characterization and the pipeline for analyzing the interactions with the environment. A) Global view of the instrument and a closed-up view of the light path (blue arrows) from the light source, through the sample, to the detection. B) Description of the working principle of the ILM detection with a schematic representation of the light path for the ILM (blue arrows) compare to the light path for the NTA (red arrows) C) The size measurement assignment of the particle is based on the analysis of its Brownian motion directly linked to the hydrodynamic radius through the Stokes-Einstein equation, where D is the diffusion coefficient, k_B_ the Boltzmann constant, T_meas_ the temperature (measurable parameter), η the dynamic viscosity and R_h_ the hydrodynamic radius. D) The diffusion of the particle, i.e. its means square displacement as a function of time lag is directly related to the diffusion mode of the EV which could be superdiffusive, diffusive or subdiffusive depending on the alpha value. In the case of particle with a known size, the measurement of the mean square displacement is then a way to access the local viscosity seen by the particles by adjusting.

In this article, we introduce an innovative approach relying on the use of interferometric light microscopy (ILM), initially developed to characterize size and concentration of (bio)nanoparticles in aqueous medium of known viscosity [30], for the local measurement of viscosity at the nanometric scale. The analysis of interference patterns allows the tracking of particles Brownian motions, which then give access to their trajectories, mean square displacement (MSD), diffusion coefficients and hydrodynamic diameters for a better comprehension of the physical interactions between particles and their environment. Contrary to other characterizations systems relying on detection inside a microfluidic chamber with injection of the sample through milli- or microfluidics, ILM detection is performed by simple deposition of the sample on a glass slide. The system of ILM is then able to work not only in aqueous solutions but also in challenging media with relatively high viscosity. Particles tracking enable the analysis of the transport properties of each nanoparticle and a global understanding of the interactions interfering with diffusion. When working with viscous matrices of unknown mechanical properties, addition of calibration steps performed with synthetic beads of various known sizes allowed the measurement of a local viscosity. Indeed, measuring the step required to match the size distribution of particles in the viscous media to that obtained in water allows for an estimation of the local viscosity. Applying this protocol to EVs encapsulated in a poloxamer, we successfully obtained information about the local viscosity at the surface of the biological vesicles and therefore allowed to quantify the impact of EV embedding in a complex matrix on their size and concentration.

## Materials and Methods

### Chemical

Phosphate-buffered saline (PBS), glycerol (reference 8.18709, CAS number 56-81-5) and poloxamer 407, named also pluronic F-127 (reference P2443, CAS number 9003-11-6), were obtained from Sigma-Aldrich (St-Louis, MO, USA). Glycerol was diluted at different concentrations (7.5%, 15% and 40%) in milliQ Water. Poloxamer 407 was diluted in PBS to achieve the expected concentrations (0.94%, 3.75%, 7.5%).

### Polystyrene (PS) beads

Beads were selected from the 3000 Series Nanosphere™ Size Standards from Thermofisher Scientific (Waltham, MA, USA), meticulously calibrated by the supplier within nanometer-scale dimensions using the trackable National Institute of Standards and Technology (NIST) methodology. They have a density of 1.05 g/cm3 and an index of refraction of 1.59 at 589 nm (25°C). PS beads stock concentration was retrieved from information given by the supplier.

### Cell culture and EVs production

EVs were produced from three cell types: Hela WT, HeLa NLuc-HSP70 [31] and human adipose-derived stem cells (hADSC) either in starvation or in turbulence (proprietary technology, WO2019002608). Cells were cultured in complete DMEM medium, i.e. DMEM (Gibco, Life Technologies Corporation, U.S.A) in which has been added 10% FBS (Corning) and 1% of penicillin / streptomycin (Gibco) at 37 °C with 5% of CO2. Geneticin (selection antibiotic) was added to complete medium at a concentration of 0,5 mg/mL for HeLa NLuc-HSP70 cells only. All cell lines were cultivated in T150 flasks, and subcultured when reaching 80% of confluency. For starvation and turbulence EV production process, EV production was performed in starvation medium (no serum) after 3 rinsing steps with PBS. For the starvation production, the conditioned medium was collected after 48h of production in T50 flasks. For the turbulence production, the conditioned medium was collected after 4h of production in 500 mL spinner-flasks. The conditioned medium was retrieved in 50 mL Falcons for clarification step using sequential centrifugations at room temperature: the supernatant was retrieved after a run at 300 g for 5 min and again after a run at 2000 g for 10 min (Eppendorf 5702). The EVs were then purified by ultracentrifugation: the pellet was retrieved after a run at 100,000 g for 90 min (Beckman Coulter Max XP, rotor MLA-50) and resuspended in low volume (~150µL) of PBS 1X. The EV-enriched fraction was stored at −80°C prior characterization using Videodrop (Myriade).

### ILM measurement

ILM measurements were performed using Videodrop instrument [32] (Myriade, Paris). Measurements were performed following the protocol from the supplier. The sample chip was washed with ethanol and / or distilled water before and after each measurement. A volume of 7 µL was deposited on the center of the sample chip for each sample measurement. The chip was then loaded on the optical path. Given the significant impact of temperature on measurements, temperature was measured for each measurement and adjusted in the software for integration into the size distribution analysis. Recordings were performed using the software version 2.5.5.6797, with a minimum of 300 particles followed per video. We exported systematically experiment summary in .pdf. and raw data in .qvir and .csv for data analysis.

### Data analysis

The analysis of data was performed on Matlab (R2021a). For each individual object, knowing it x and y position over time, the trajectory coordinates were retrieved from the csv files. The MSD was then calculated using (Eq. 2), with an average along the trajectory. The fitting of the MSD was performed using the function for nonlinear fitting ‘fitnlm’ and performed only on the first 10 points of the MSDs. Indeed, the trajectory is given for a maximum of 100 points (if the particle was followed over the whole film) and a minimum of 10 points. For each value over time, the MSD corresponds to the average of the MSD value obtained for all the segments of *nδt* with *n* = [1; 100]. Therefore, the average is made on a decreasing number of segments over time, resulting in a very noisy curve for late time points. In consequence, the fitting step was performed systematically on the first 10 points to optimize fitting accuracy regardless of trajectory durations.

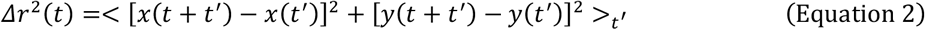

For each figure involving MSD, we plotted the geometric average of the MSD over the N (>300) particles followed. The plots were made by using the function ‘shadedErrorBar’ [33] to display the standard error of the mean (SEM) and the function ‘al_goodplot’ [34].

### Macroscopic viscosity measurement

Macroscopic viscosity measurements were performed using a Physica RheoCompass MCR 302 (Anton Paar, France) equipped with a solvent trap for preventing solvent evaporation. The measurements were performed using a cone and plate geometry (diameter = 50 mm, cone angle = 1°). Temperature was controlled during measurement using a Peltier plate unit adjusted at 24°C and was regulated during 3 min prior each measurement. Viscosities were measured using a shear gradient of 100 Hz during 3 min at 0.94%, 1.87%, 3.75%, 7.5%, 20% and 30%.

### Electron microscopy imaging

Grids for electron microscopy were prepared using an automated plunge freezer (EM-GP, Leica). 3.5 µL of samples were deposited on a glow discharged lacey holey carbon grids (Ted Pella INC., Lacey grids) equilibrated at 10°C. Samples were blotted during 3-5 sec to remove the excess sample and leave a thin film in the carbon hole. The blotting was carried out on the opposite side from the liquid drop and plunge-frozen in liquid ethane at −181 °C. For high viscosity samples, an equilibration time of 10 to 15 seconds was added and blotting time was adapted to the viscosity of the sample, from 3 to 20 seconds. The samples were observed at liquid nitrogen temperature using a Tecnai F20 (Thermofischer, FEI, Eindhoven, the Netherlands) microscope operated at 200 kV and equipped with a Falcon II direct electron detector (Thermofischer, FEI, Eindhoven, the Netherlands). The data were collected at a magnification of 50 000 for a pixel size of 2Å per pixel.

## Results

### Part I Characterization of the ILM instrument and nanorheology

Imaging and characterizing nanoparticles remains a significant challenge in the field of nanotechnology. Optical imaging, renowned for its non-destructive, robust, fast, and user-friendly nature, emerges as a primary technique for this purpose. Nonetheless, as nanoparticles diminish in size (< 200 nm) and breach the diffraction limit, optical methods alone struggle to effectively visualize and characterize them. To address this limitation, several alternative methods have been proposed, among which Dynamic Light Scattering (DLS), Nanoparticle Tracking Analysis (NTA) and (nano)flow cytometry stand out. These techniques rely on the detection of the scattering signal of nanoparticles at an angle of 90° of the light path, which, according to the Mie theory, is proportional to *R*_*h*_^6^ (with the assumption that the wavelength of the light source λ>> R_h_). Consequently, as the size of nanoparticles decreases, there is a significant loss of signal. Thus, despite advancements, distinguishing nanoparticles from the noise background in images remains a formidable challenge.

To address this challenge, the interferometric light microscopy (ILM) detection relies on the detection of the signal from nanoparticles directly on the optical path (180°), taking advantage of the interferences between the scattered and the LED source light. The intensity of the interferences between the light source and the scattering light from the particles on the optical path can be described as:

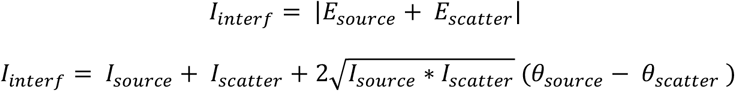

With *I*_*source*_ the light source intensity, *I*_*scatter*_ the intensity of the scattering light from the particles and *θ* the phase of each light source.

As described in Figure 1 A and B, the light source, sample and camera detector are aligned for the detection of interference patterns. The signal strength is then proportional to the *R*_*h*_^3^, mitigating the signal weakness associated with small particles.

The ILM technique enables the detection of nanoparticles within a liquid matrix, as presented in Figure 2 B. Custom image processing, developed by Myriade, is employed on each image to isolate and subtract the static signal emanating from the LED, from the dynamic signal generated by spontaneous particle movements due to their Brownian motion (Figure 2A). A distinct pattern made of a white and black doublet represent interference patterns produced by nanoparticles (Figure 2A), whose position can then be precisely extracted over time. Particles must stay in focus long enough (at least 10 successive images) to be able to be track by the system, and thus analyzed for size quantification. If it is not the case, particles are not considered for diffusion measurement and size quantification. These two kinds of particles are identified by with orange and white circles respectively on Figure 2B.

**Figure 2:**
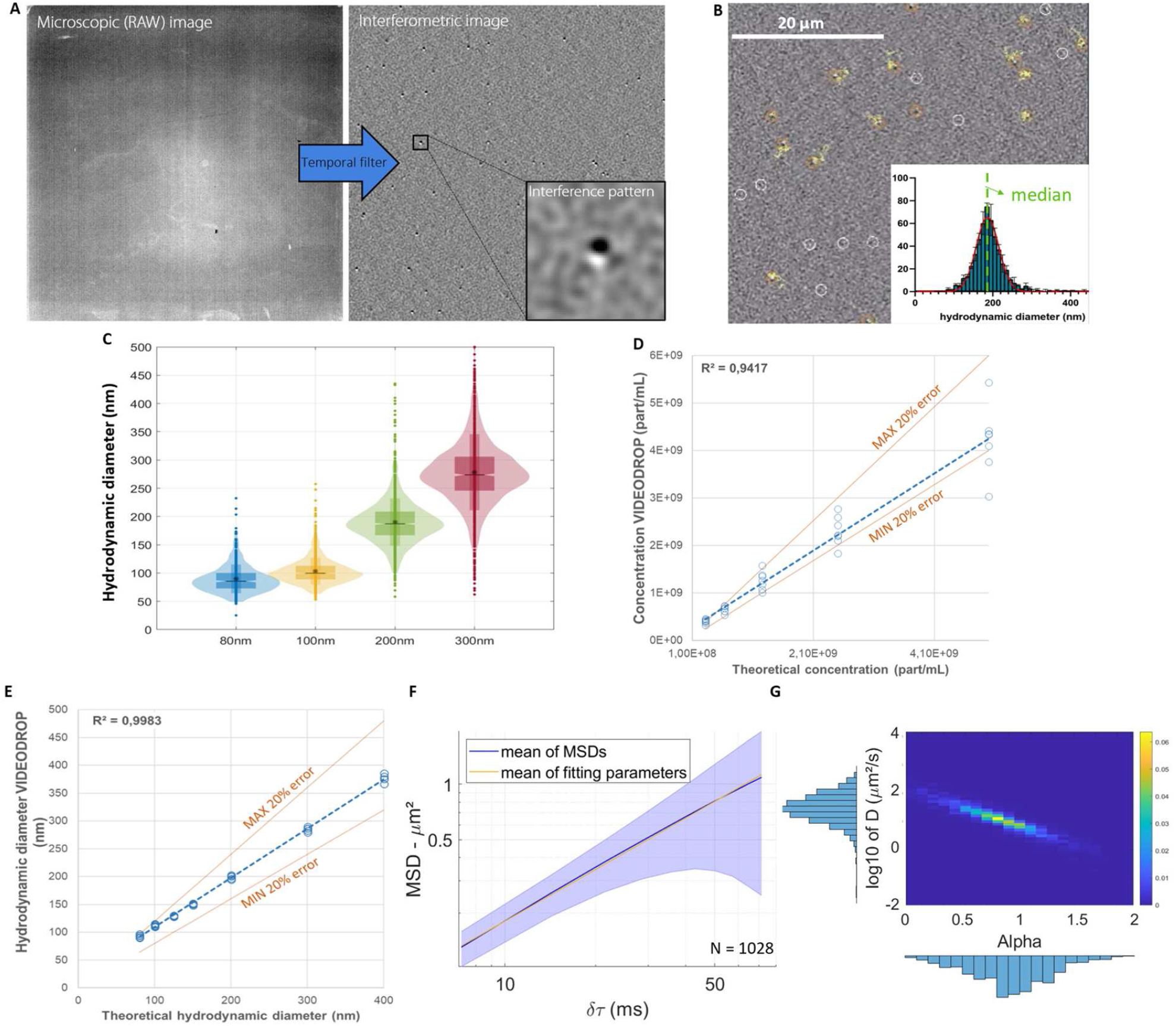
Description and characterization of the ILM instrument for PS beads quantification in water. A) Interferometric image generated from the microscopic one using temporal filtering and allowing to visualize interferometric pattern related to nanoparticle. B) Screen capture of the particle visualization by ILM and presentation of the results by the instrument. C) Presentation of size distributions of the different size of beads. Description of the linearity of the measure of D) concentration E) size. F) Mean MSDs of 100nm beads in water with the SEM distribution around it and the equation made of the mean of each parameters G) Distribution of diffusion coefficients (D) and the anomalous diffusion exponent (α) extracted from each measured particle

By conducting measurement within a predetermined volume of the sample, this method provides direct access to the concentration. Moreover, if working with a sample of known or pre-calibrated viscosity, the size distribution can be inferred from the Brownian motion recorded for each particle, giving access to the particle diffusion. Particle diffusion coefficient is correlated with size through the Stokes-Einstein equation (Equation 1), as presented on Figure 1 C.

To leverage this property of the ILM for hydrodynamic size assessment, we initially employed PS beads that were meticulously calibrated by the manufacturer. These beads serve as exemplary standards with precisely known sizes. Dilution in PBS was performed for each particle size to achieve the instrument’s measured range, following which they were imaged (Figure 2B) to assess their size distribution (Figure 2C).

We evaluated the linearity of nanoparticle concentration and size measurements, demonstrating their consistency across a concentration range from 10^8^ to 10^10^ part/mL (Figure 2D) and a hydrodynamic diameter range of 80 to 400 nm (Figure 2E). Notably, accuracy remained within 10% for sizes ranging from 200 to 400 nm. Further analyses were performed with beads of 100, 200 and 300 nm in diameter, mimicking biological samples of interest.

However, all these assertions can only be reliably made under the condition of purely diffusive particles in a uniform viscous medium with a low Reynolds number. To ensure that these conditions are met, the MSD of each particle can be measured, as depicted in Figure 2F, to detect any restriction of movement (subdiffusive trajectories) or acceleration of movement (superdiffusive trajectories). Each of the MSD can be fit by a power law following the form:

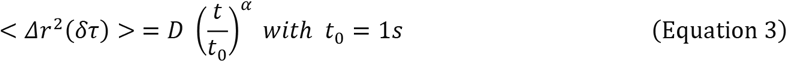

Where *r*^2^ is the space explored by the particle during an interval of time *δτ, D* is the diffusion coefficient and *α* is the anomalous diffusion exponent. A diffusive state is characterized by a time coefficient α= 1; if α > 1, the movement is superdiffusive, if α< 1, the movement is subdiffusive (Figure 1D).

The displacement of each particle allows the calculation of the MSD of the recorded trajectory for each particle. The 10 first points of the MSD curve were then fitted using a power law. The mean of the MSDs is depicted in blue on Figure 2F, along with the standard error of the mean (SEM) around it. In yellow, the power law corresponding to the geometric mean of diffusion coefficients (*D* = 0.0213 + 0.0013/ −0.0004 µ*m*^2^/*s*) and arithmetic mean of time coefficients (*α* = 0.93 ± 0.01) is represented. The diffusion and time coefficients of each particle are illustrated in Figure 2G and Supplementary Figure 1. A correlation between the diffusion and time coefficients was observed for particles of all three sizes. Indeed, we observed a decrease in *D* while *α* decreases. This correlation could be related to the disparity of the MSDs around the diffusion, as presented on Supplementary Figure 2.

In the case of diffusive motion of particles at a constant temperature, the size measurement becomes dependent solely on the dynamic viscosity of the liquid phase surrounding the particles. This nanoscopic viscosity represents the local viscosity experienced by the measured particles. While the Videodrop instrument can be used to determine the hydrodynamic size for a known viscosity (usual application of this instrument), it can also be used to determine the viscosity for a known hydrodynamic size. Indeed, when measuring the size distribution of particles of calibrated size in a viscous liquid of unknown viscosity, a shift in the size distribution occurs. This shift is then compensated for by accounting for the viscosity. The viscosity value can be adjusted in order to match the median of calibrated nanoparticles (Figure 1D). This step relies on a precise measurement of the size distribution of calibrated particles as presented on Figure 2 C and will be develop in the second part of this paper.

### Part II Matrix viscosity measurement using calibrated particles (measurement of *η* with *R*_*h*_ known)

To investigate the diffusion of nano-objects in a complex environment, it is crucial to be able to measure the local viscosity of the complex material. In this part, we propose to measure this viscosity on bead with a known and well characterized size (known *R*_*h*_). As a consequence, the only unknown parameter in the Stokes-Einstein equation (Eq. 1) is the viscosity 1 of the matrix. We used this approach in two different media: glycerol, a Newtonian fluid, and poloxamer 407, a non-Newtonian matrix.

First, we conducted ILM measurements of calibrated PS beads in a Newtonian fluid, specifically glycerol at different glycerol concentrations (7.5, 15, 40%), corresponding to various macroscopic and nanoscopic viscosities of the fluid. Size distributions were obtained by first setting the viscosity to the one of water (1 mPa.s). Viscosity was then progressively adjusted until the median hydrodynamic diameter matches the theoretical diameter of the calibrated particles. The viscosities obtained with the three particle sizes across different glycerol concentration gave very similar values (Figure 3C), with discrepancies emerging particularly at higher glycerol concentrations. These variations in measurements may be attributed to an accelerated drying process at high glycerol concentrations. The size distributions of beads (diameters of 100, 200, and 300 nm) obtained with the final value of viscosity are depicted in Figure 3A. The size distribution appears to be more spread, particularly for larger beads, at higher concentrations of glycerol. This could be explained by a decrease of the localization precision of the bead at higher hydrodynamic diameter and higher viscosity. Indeed, an increase in viscosity and particle size leads to a decrease in the displacement observed between two temporal points. And the smaller the displacement is, the higher the imprecision on the trajectory point is.

**Figure 3:**
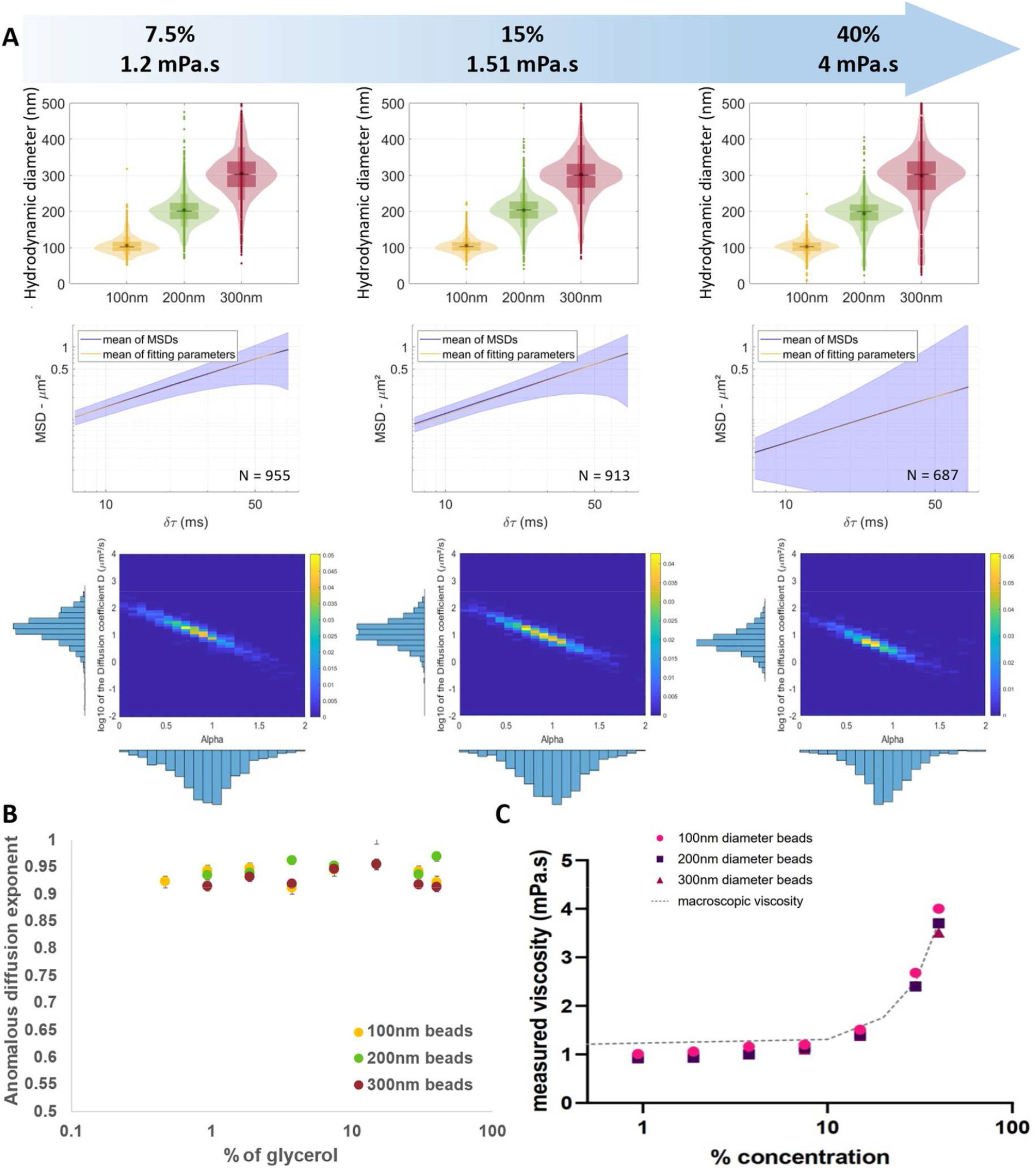
Validation of ILM as a nano-rheometer in a Newtonian fluid, the glycerol. A) representation of the size distribution of 100, 200 and 300nm diameter beads with the mean MSD (+/− sem) of particles and the relation between the diffusion and time coefficients of the MSD profile of 100nm beads in glycerol at 7.5, 15 and 40%, showing a diffusive pattern, and B) Representation of the distribution of the time coefficient alpha for 100nm, 200nm and 300nm diameter beads in 40% glycerol ±SEM C) measured viscosity at the surface of the beads in different concentrations of glycerol

To ensure the diffusive nature of particle trajectories within the glycerol matrix, we computed the MSD of each particle and fitted them with a power law, as previously described. Hundreds of particles were analyzed for each condition. The mean of the MSDs is depicted in blue, along with the standard error of the mean (SEM). Additionally, in yellow, the power law with parameters corresponding to geometric mean of diffusion coefficients (*D* = 0.0055 + 0.0003/−0.0001µ*m*^2^/*s* for 40% of glycerol) and arithmetic mean of anomalous diffusion exponents (*α* = 0.92 ± 0.01 for 40% of glycerol) is presented in Figure 3A. Remarkably, for all tested glycerol concentrations, no difference was observed between the two curves, indicating the efficacy of our fitting method, as demonstrated in Supplementary Figure 3.

For each particle, we were able to extract a diffusion coefficient (*D*) and an anomalous diffusion exponent (*α*) from the MSD power law fitting. The heatmap of the diffusion coefficients as a function of the anomalous diffusion exponents is presented in Figure 3A. This graph revealed a correlation between the two coefficients, similar to observations in water. The anomalous diffusion exponent *α* is consistently centered around 1, meaning that particles are exhibiting diffusive behaviors by freely diffusing in the viscous complex environment. The diffusion coefficients were centered around 10 µm^2^/s, a value consistent with findings in the literature [35,36]. Additionally, the diffusion coefficient demonstrated a decrease with increasing particle size and glycerol concentration, as depicted in Supplementary Figure 4. In these conditions, the anomalous diffusion coefficients (*α*) remained centered around 1 for beads of all three sizes and across all concentrations of glycerol, as illustrated in Figure 3B.

The nanoscopic viscosity measured by ILM for each glycerol concentration was then compared to the macroscopic viscosity, as depicted in Figure 3C. The good agreement between the nanoscopic and the macroscopic viscosity successfully validated our viscosity measurement method and protocol and is in agreement with the Newtonian nature of glycerol.

In the following, we will work on a more complex and biologically relevant material, the poloxamer 407. We propose to use the same methodology to determine the viscosity of this material.

The same PS beads as the one used previously were tracked within a non-Newtonian matrix, poloxamer 407 to determine its nanoscopic viscosity. We worked with different concentrations of the poloxamer (0.94%, 3.75%, 7.5%) corresponding to different organization of the matrix but always in liquid phase of various viscosities. We used the same method as above, to deduce the viscosity of the poloxamer from size distribution. This bead size distribution was assessed across the three concentrations of poloxamer, unveiling a broadening of the size distribution at higher concentrations of poloxamer. This phenomenon was similarly observed with increasing concentration (and therefore viscosity) of glycerol (Figure 4A). For the 300 nm beads, the emergence of an artefactual subpopulation of smaller particles was observed.

**Figure 4:**
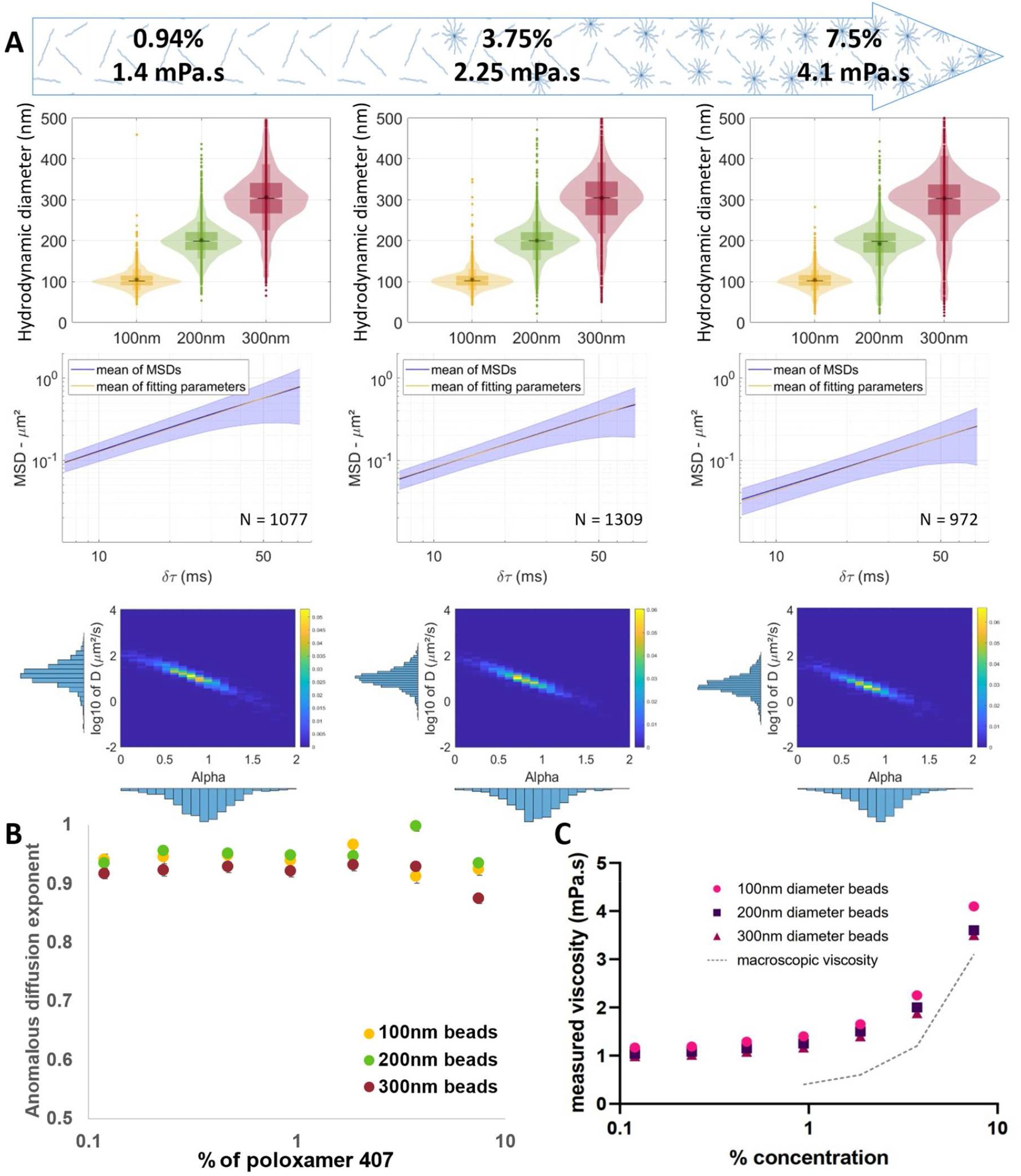
Measurement in a non-Newtonian fluid the poloxamer 407. A) representation of the size distribution of 100, 200 and 300nm diameter beads with the mean MSD of particles and the relation between the diffusion and time coefficients of the MSD profile of 100nm beads in poloxamer 407 at 0.94, 3.75 and 7,5%, showing a diffusive pattern, and B) Representation of the distribution of the anomalous diffusion exponent for 100nm diameter beads in 7,5% poloxamer 407 ±SEM C) measured viscosity at the surface of the beads in different concentrations of poloxamer 407

MSDs, diffusion coefficients, and anomalous diffusion exponent were computed for the 3 sizes of PS beads across various concentrations of poloxamer. The mean MSDs remained congruent with the function derived from the mean of each power law parameters (*D* = 0.0051 + 0.0006/−0.0001 µ*m*^2^/*s* and *α* = 0.92 ± 0.01 for 7.5% of poloxamer), similar to observations in water and glycerol, validating here our fitting method in a non-Newtonian matrix. The diffusion coefficient exhibited a decrease as bead size or poloxamer concentration increased (supplementary Figure 6), with values in agreement with the literature [37]. Conversely, the anomalous diffusion exponent *α* consistently remained centered around 1 (Figure 4B), indicative of a pure diffusive pattern (pure Brownian motion) even in this complex matrix, allowing to extract a measure of the viscosity and confirming the validity of the Stokes-Einstein equation for this matrix.

Interestingly, the nanometric viscosity quantified by using Videodrop as nanorheometer, exhibited a noticeable disparity compared to the macroscopic viscosity (Figure 4.C). Several factors could account for this observation. Firstly, it is important to consider the non-Newtonian behavior of poloxamer 407, which is a non-homogeneous medium in which nanoparticles probe the local environment at their scale, i.e. the nanoscale. These interactions may impose restrictions on bead diffusion within the poloxamer matrix, potentially leading to complex non-linear diffusion patterns. One such model that could be applicable is the Hopping diffusion model, which accounts for non-linear behavior in diffusion processes. Furthermore, it is crucial to note that in non-Newtonian fluids, the viscosity is dependent on the scale at which it was measured. Therefore, the macroscopic viscosity, which we typically measured and observe on a larger scale, may not accurately represent the nanoscopic viscosity experienced by individual particles measured by ILM. The intricate structure and behavior of non-Newtonian fluids can result in significant disparities between macroscopic and nanoscopic viscosities, further complicating our understanding of particle dynamics within these systems. Nevertheless, we found that ILM is a good methodology for assessing rapidly the nanoscopic viscosity of poloxamer and we are confident that this methodology could be used on other complex (non-)Newtonian media.

### Part III Size distribution measurement from biological samples inside characterized matrix (measurement of R_h_ with η known)

In the previous section, we successfully determined the local nanoscopic viscosity of complex matrices by working with calibrated particles. Leveraging this measured parameter, we now investigate the diffusion patterns of bionanoparticles such as EVs in complex environment. EVs are highly heterogeneous in size and our first step consists in assessing the instrument’s measurement capabilities on a controlled polydisperse sample. Measurements were performed on mixes of PS beads in an aqueous fluid (water-like viscosity): phosphate-buffered saline (PBS) (Figure 5.). For each mix, the obtained size distribution (Figure 5B-C) was compared to the simulated one corresponding to the linear combination (Equation 4) of the monodisperse bead size distribution (Figure 5.A) weighted by the mixture rates.

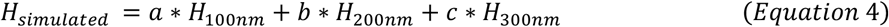

**Figure 5:**
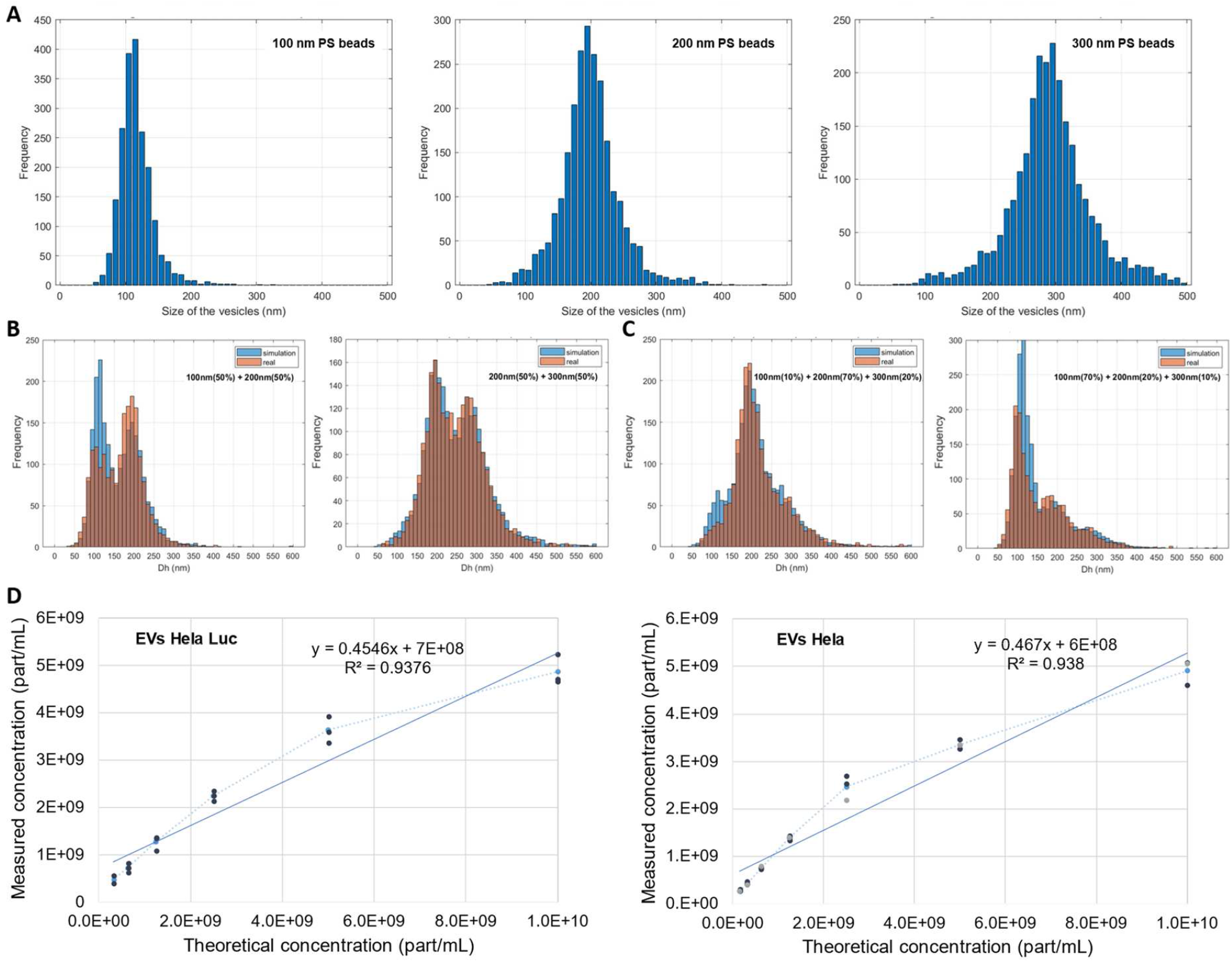
Validation of ILM for polydisperse samples characterization. A) Size distribution of monodisperse PS beads of 100nm, 200nm and 300nm beads. B) Simulated and measured size distribution of a binary mixtures of 100 and 200 nm (left, with a=0.5, b=0.5, c=0) beads and 200 and 300nm beads (right, with a=0, b=0.5, c=0.5). C) Simulated and measured size distribution of ternary 100nm/200nm/300nm beads mixtures (left: a=0.1, b=0.7, c=0.2, right: a=0.7, b=0.2, c=0.1). D) Linearity of the concentration measurement of EVs from two Hela cell lines (WT and HSP70-Luc).

Where a, b and c denote the respective percentages of beads of 100 nm, 200 nm and 300 nm size respectively. These comparisons demonstrate the high efficacy of the system in detecting subpopulations of beads within mixtures comprising two or three distinct populations, with precise identification of each peak. This property is particularly crucial when studying EVs, given their inherent polydispersity in size.

In a second step, we validated the use of ILM for EV concentration in aqueous solution, at different concentrations for EV from two different cell lines. High degree of linearity for naturally polydisperse biological sample was observed across a broad concentration range from 10^9^ part/mL to 5.10^10^ part/mL for EVs in PBS (Figure 5D). The same range was identified for EVs derived from two distinct cell lines.

In the third step, we combined both usages of ILM: (i) *nanometric viscosity quantification* based on ILM analysis of particles of known size in a medium (non-Newtonian poloxamer) of unknown viscosity, (ii) *size quantification* based on ILM analysis of particles of unknown size in a medium of known viscosity. Indeed, we conduct ILM measurements of EVs in poloxamer 407, which viscosity was measured with ILM above (Part II).

As previously, we observed a good agreement between the geometric average of MSDs and the power low resulting from the average fitting parameters (Figure 6A). For both cell types, EVs behave as diffusive particles with *D* = 0.0083 + 0.0003/−0.0001 µ*m*^2^/*s* and *α* = 0.91 ± 0.01 for HeLa-derived EVs and *D* = 0.0097 + 0.0004/−0.0001 µ*m*^2^/*s* and *α* = 0.88 ± 0.01 for hADSC-derived EVs. The heatmaps corresponding to EVs in PBS (Figure 6A) closely resembled to those of beads in water (Figure 2F,G), with an anomalous diffusion exponent centered around 1, in agreement with a purely diffusive regime. However, when embedded in poloxamer, EVs diffusion is impacted: both the diffusion coefficient decreases and the anomalous diffusion exponent exhibits values lower than 1 (*D* = 0.0028 + 0.0002/−0.0001 µ*m*^2^/*s* and *α*= 0.78 ± 0.01 for HeLa-derived EVs and *D* = 0.0090 + 0.0005/−0.0001 µ*m*^2^/*s* and *α* = 0.93 ± 0.01 for hADSC-derived EVs. Therefore, EVs in poloxamer exhibit motions corresponding to subdiffusive regime. The observed decrease in both anomalous diffusion exponents and diffusion coefficients suggests an interaction between poloxamer and EVs, resulting in restricted EV movement. This phenomenon is supported by the characteristic shape of the MSDs, exhibiting periodic patterns indicative of a jerky trajectory (supplementary Figure 5).

**Figure 6:**
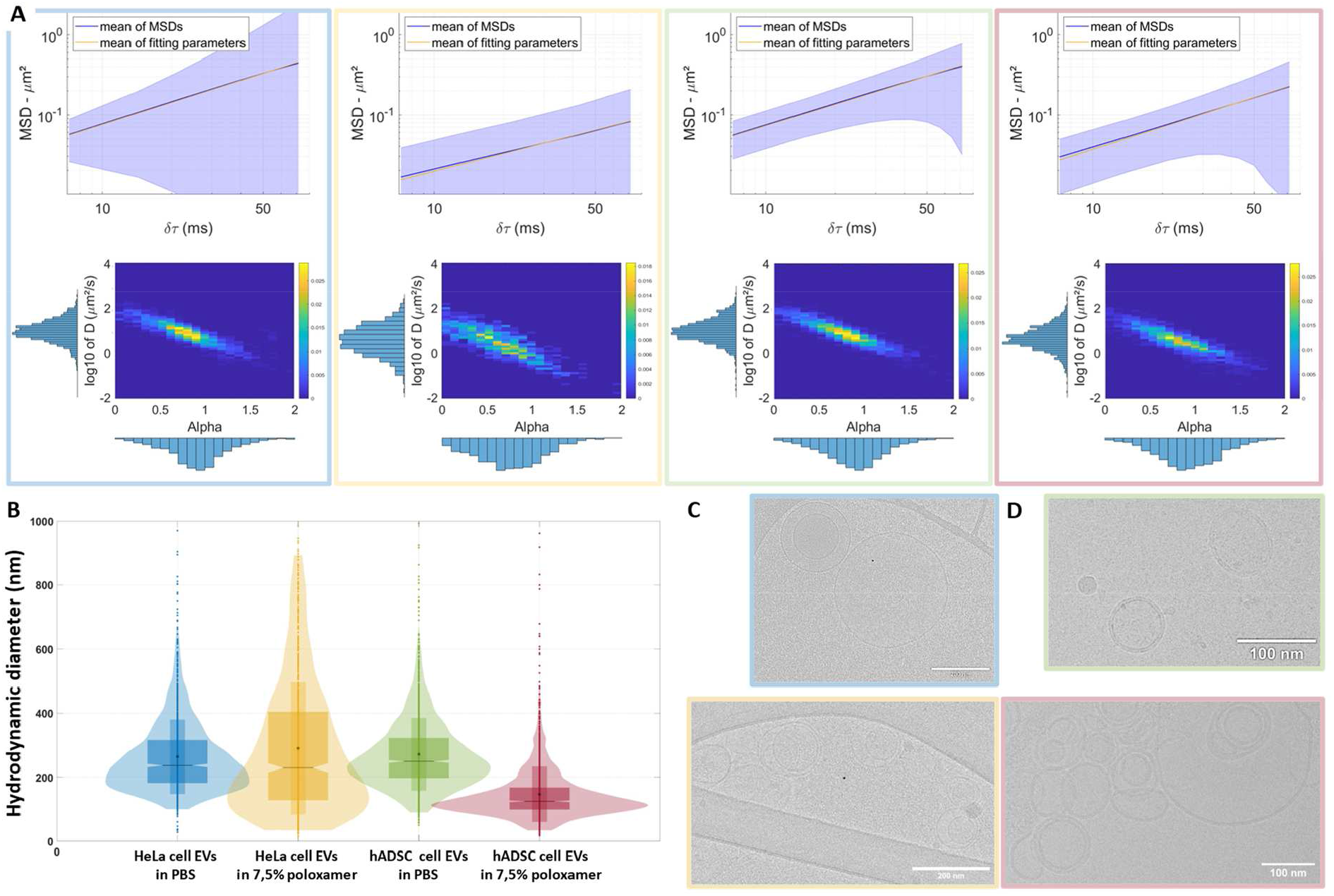
Characterization of EVs by ILM. A) representation of the MSD, the diffusion and time coefficient of EVs from Hela and hADSC cells in PBS and in 7,5% of poloxamer 407 B) Display of the size distribution of EVs from the two previously introduced cell lines measured by ILM on PBS and in 7,5% of poloxamer 407 C) images by cryoEM of EVs from Hela cells in PBS and 7,5% of poloxamer 407. D) images by cryoEM of EVs from hADSC cells in PBS and 7,5% of poloxamer 407

The spread size distribution of EVs observed in PBS demonstrates the heterogeneity of EVs in terms of size. EVs derived from Hela and hADSC cells exhibited relatively similar size distributions in PBS. Interestingly, both the concentration and size of EVs decreased when measured in poloxamer for both cell types. However, the poloxamer affected differently for each cell type the size distribution shape: Hela cell-derived EVs exhibited an enlargement in distribution, while hADSC cell-derived EVs showed a sharpening of size distribution, indicating an inverse effect.

CryoEM enabled direct visualization of EVs in both PBS and poloxamer matrices. In both environments, the bilayer structure of EVs was clearly observed, demonstrating the presence of intact EVs. In poloxamer, however, we observed evidence of tension on some EVs, with apparition of broken membranes and release of proteins, explaining the apparition of new nanometric entities affecting size distributions.

## Discussion

While the use of EVs in nanomedicine is rapidly expending, there is an urgent need of developing new technologies for characterizing EVs in complex media involved in innovative delivery systems. In this frame, our team as focused on the use of poloxamer 407 (Pluronic F-127), a thermosensitive gel (liquid at room temperature and jellified at body temperature), particularly interesting for the treatment of fistula [27] To transit to the gel phase at body temperature, a concentration of 20% of poloxamer was found to be optimal. However, at this concentration at room temperature, the viscosity is around 400 mPa.s (400 times the viscosity of water). An attempt to characterize EVs in a 20% poloxamer fluid using NTA (LM 10, Malvern Panalytical) was performed in this previous quantitative study of our group, however the displacement of particle due to the Brownian motion was difficult to observe, rending impossible the trajectory analysis and therefore the retrieval of hydrodynamic radius. At the best, a final concentration of poloxamer at 2% has given analyzable data. Using an alternative technology to NTA, namely interferometric light microscopy (ILM), we are able in the present work to characterize poloxamer in concentrations up to 7.5%. Macroscopic viscosity of poloxamer 407 at this concentration at room temperature can be measure using standard rheology. However, as poloxamer is a non-Newtonian fluid, we wanted to use viscosity that is measured at the same scale as the target object: EVs. In this work, we develop an innovative technique using ILM to measure the viscosity of (complex) fluid at the nanoscale by relying on the Stokes-Einstein equation. Indeed, in a diffusive regime, by analyzing the nanoparticles trajectory due to their spontaneous motions, the Stokes-Einstein equation allow us to relate the viscosity, the hydrodynamic size and the temperature. The temperature being a measurable parameter, (i) if the viscosity is known we can deduce the hydrodynamic size from the Brownian motion (major usage of this method), (ii) reciprocally, if the size of the particles is known, the viscosity can be deduced. After a meticulous analysis of the performance of the ILM system to characterize properly synthetic monodisperse nanoparticles of various size, we tested the ability of the system to assess properly the size and concentration of polydisperse sample (mix of various sizes of synthetic beads ranging from 100 to 400 nm or EV sample). In each case, we ascertain the range within which the system was subjected to linear concentration. Our approach for using Videodrop instrument as nano-rheometer, involved the use of calibrated nanoparticles to assess the nanoscopic viscosity of the surrounding matrix. This method was tested on both Newtonian and non-Newtonian fluids. For Newtonian fluids, the nanoscopic viscosity values were in agreement with the macroscopic values, whereas for non-Newtonian fluids, nanoscopic measurements yielded slightly higher values compared to macroscopic assessments. Then, we leveraged these viscosity measurements to analyze the size distribution of biological nano-objects within a non-Newtonian complex matrix. It is important to emphasize that the measured viscosity represents a nanoscopic local viscosity that is precisely experienced by the nanoparticles. This approach was applied to the size measurement of biological samples of EVs within environment characterized at the nanoscale and with the same instrument. When examining EVs in the poloxamer, we observed distinct forces acting on the biological particles that could lead to size distribution distortion. The matrix seems to constraint the movement of EVs. Additionally, as EVs are deformable nano-objects, their size and shape could be potentially altered by the poloxamer, as it was suggested by cryoEM images. Furthermore, the release of proteins from EVs may contribute to changes the local viscosity of the poloxamer, impacting even further the transport behavior of both EVs and other particles within the matrix. By comparing EVs produced from two different cell types, we highlighted that the effect on EVs seems to be cell type dependent. Indeed, both the average hydrodynamic diameter and the size distribution shape were impacted more strongly for hADSC than for HeLa-EVs. This effect could be related to a slightly different composition of EV membrane, as it was described that poloxamer (also called Pluronic block copolymers) due to its amphiphilic properties behave like surfactant with the ability to interact with biological membranes [38]. A different impact of poloxamer on EVs could happen at higher concentration of poloxamer, as it was described in this same paper that the formation of micelles diminishes the capability of poloxamer to affect cellular membranes.

The phenomenon of hopping diffusion, characterized by non-linear movement patterns (i.e. alternance of confined and diffusive motions), could be an explanation for observed patterned trajectories of EVs in poloxamer. This behavior suggests that the transport of particles within the matrix is influenced by complex interactions between the particles and the polymer network. Understanding these interactions is crucial for accurately predicting particle movement in viscoelastic environments, but asks for deeper analysis of trajectory like it was done recently in the literature [39,40].

The utilization of ILM in this study has provided valuable insights into the interplay between nanoparticle transport and nano-rheology within complex fluid. By employing ILM, we were able to not only measure nanoparticle concentration and size distribution, but also highlight interactions of nanoparticles with their environment in a comprehensive manner. This approach represents a significant advancement in the characterization of nanoparticles, particularly in complex and viscous media, which are often encountered in the context of gel-based EV formulations. The results obtained from this study underscore the importance of considering both intrinsic nanoparticle properties and the rheological characteristics of the surrounding matrix when designing targeted nanoparticle therapies for solid tumors. Furthermore, our study highlights the potential of ILM as a valuable tool for characterizing nanoparticles in complex biological environments. Unlike traditional microscopy techniques, ILM offers several advantages, including high sensitivity, real-time imaging capabilities, and the ability to work in challenging media with relatively high viscosity. This makes ILM particularly well-suited for studying the interactions between nanoparticles and their surrounding environment in biologically relevant contexts.

Overall, our findings shed light on the intricate interplay between particles and their surrounding matrix. The integration of ILM for rheological analysis represents a promising approach for studying nanoparticle transport in challenging biological environments. By elucidating the forces at play and the mechanisms governing particle transport in viscoelastic matrices, we pave the way for the development of more effective biological carriers and therapeutic interventions.

## Conclusion

In conclusion, the utilization of interferometric light microscopy (ILM) presents a promising avenue for the characterization of nanoparticles, including extracellular vesicles (EVs), especially within complex matrices of high viscosity. By combining the advantages of ILM, such as its ability to measure particle concentration, size distribution, and track particle trajectories, with calibration-like procedures and validation against known standards, we have demonstrated its efficacy in quantifying the mechanical properties of both Newtonian and non-Newtonian fluids. Furthermore, our study highlights the potential of ILM to provide insights into the interactions between nanoparticles and their surrounding environment, offering valuable information for various biomedical applications, including targeted drug delivery and tissue regeneration. Moving forward, the continued development and refinement of ILM techniques hold promises for advancing our understanding of nanoparticle physical behavior and facilitating their utilization in diverse biomedical contexts.

## Aknowledgements

The authors would like to aknowledge Gregory Lavieu for sharing HeLa cell lines (WT and Nluc-HSP70), François Mazuel and Donatien Ramiandrisoa (Myriade Lab) for their precious clarifications about QVIR software, Kondareddy Cherukula for sharing his EV productions and Gérard Pehau-Arnaudet from UBI Facility for his expertise in cryoEM. M.Nilges for purchasing the Falcon II detector financed by the Equipex CACSICE (grant number ANR-11-EQPX-0008) Videodrop instrument, equipment for EV production and purification are part of the IVETh expertise facility. IVETh is supported by the IdEx Université Paris Cité, ANR-18-IDEX-0001, by the Region Ile de France under the convention SESAME 2019 – IVETh (EX047011) and via the DIM BioConvS, by the Region Ile de France and Banque pour l’Investissement (BPI) under the convention Accompagnement et transformation des filières projet de recherche et développement N° DOS0154423/00 & DOS0154424/00, DOS0154426/00 & DOS0154427/00, and Agence Nationale de la Recherche through the program France 2030 “Integrateur biotherapie-bioproduction” (ANR-22-AIBB-0002). L. Alexandre is supported by PEPR CARN.

