## Supplementary Figures for "*In situ* investigation of extracellular vesicles in viscous formulations: interplay of nanoparticle transport and nano rheology through interferometric light microscopy analysis"


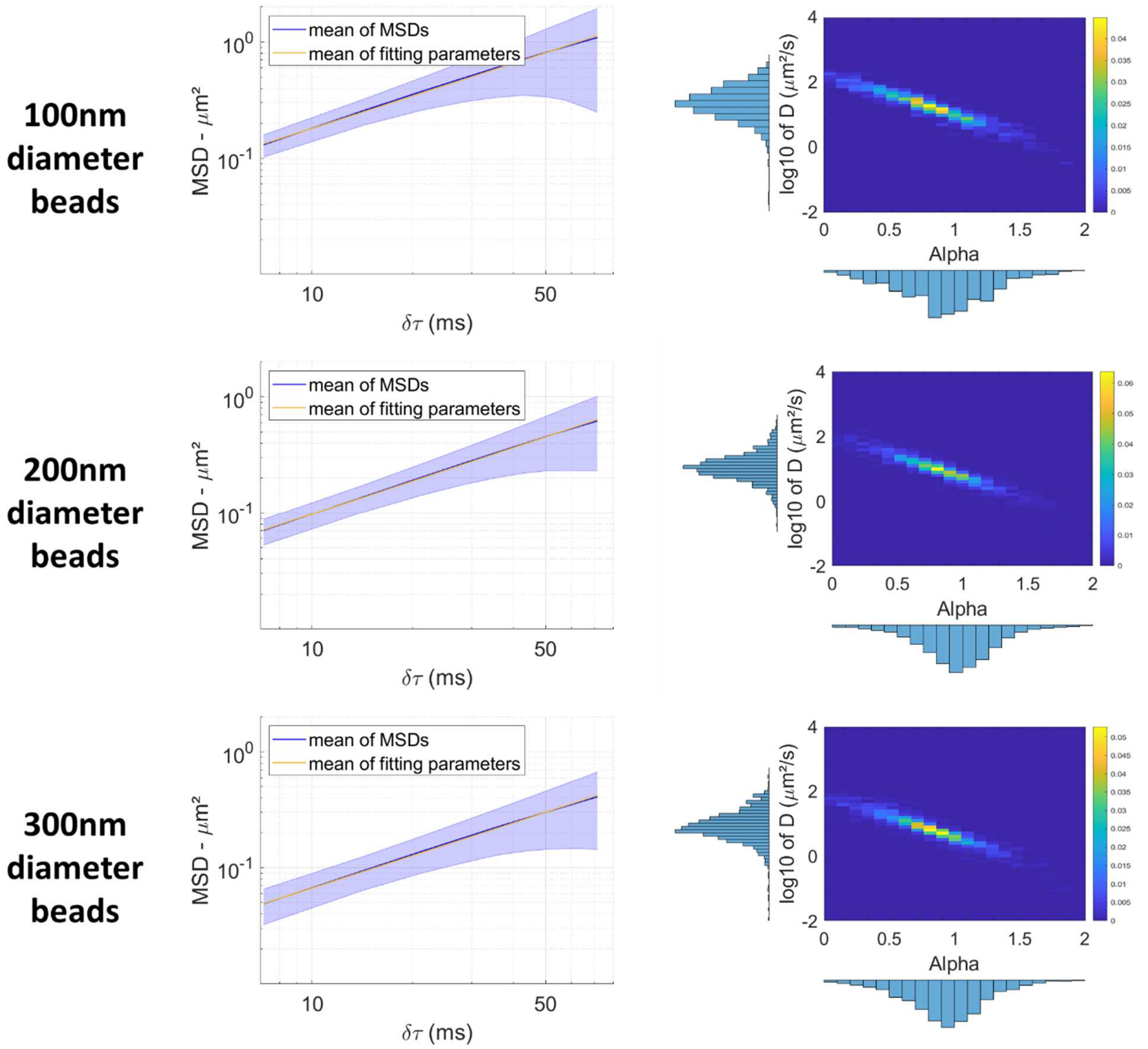


*Supplementary Figure 1: Geometric average MSD for 100 nm, 200 nm and 300 nm beads in PBS and the heatmaps of diffusion coefficients as a function of the anomalous diffusion exponents obtained for each MSD.*

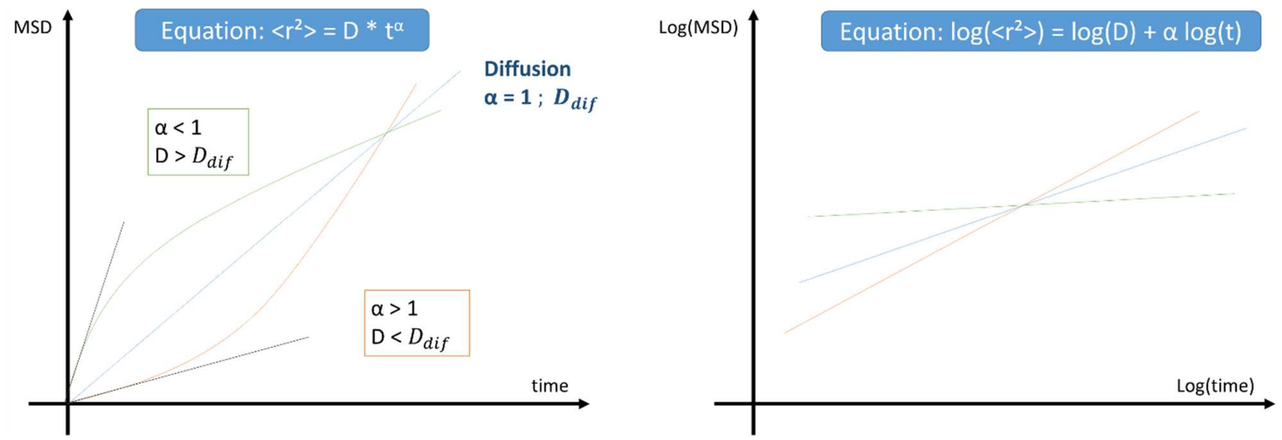


*Supplementary Figure 2: graphical simulation potentially explaining the correlation between e diffusion coefficient D and the anomalous diffusion exponent α (left: in linear scale, right: in log scale).*

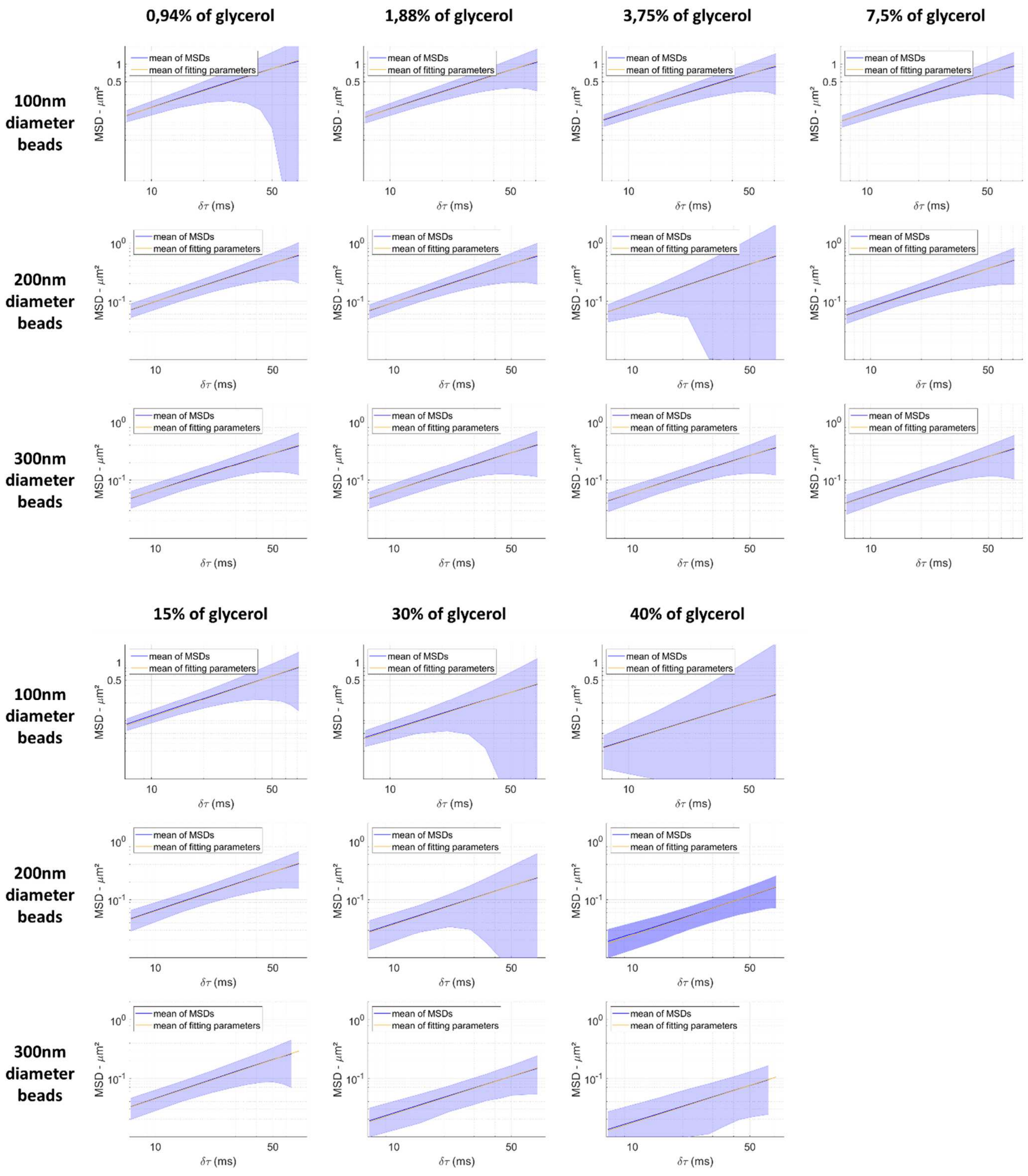


*Supplementary Figure 3: Geometric average MSD +/- SEM for 100 nm, 200 nm, 300 nm PS beads embedded in 0.94%, 1.88%, 3.75%, 7.5%, 15%, 30% and 40% of glycerol.*

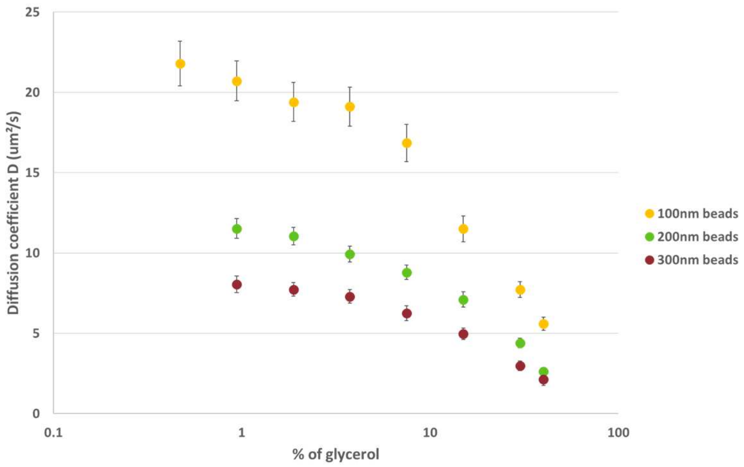


*Supplementary Figure 4: Diffusion coefficient of 100 nm, 200 nm and 300 nm PS beads as a function of the percentage of glycerol.*

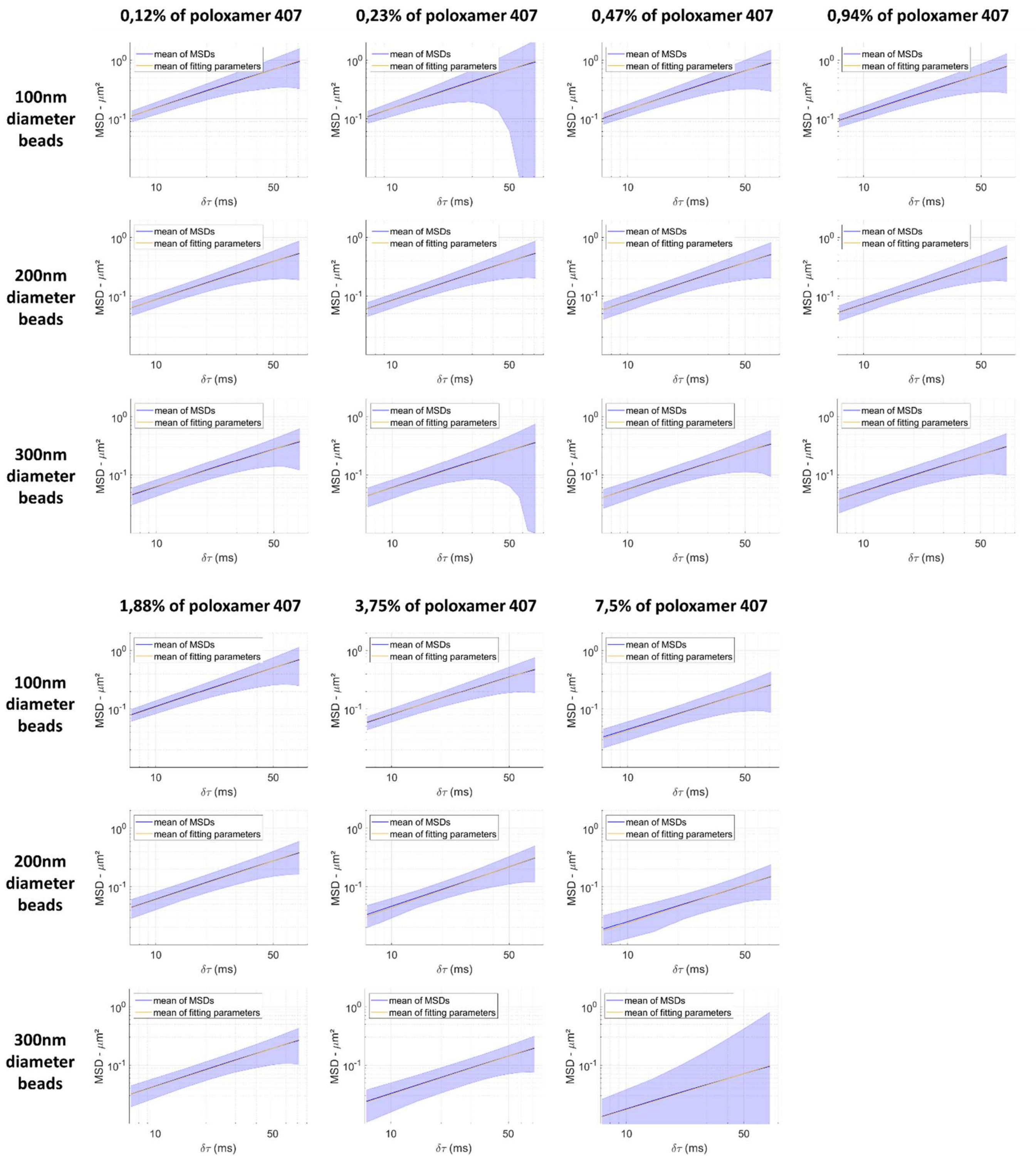


*Supplementary Figure 5: Geometric average MSD +/- SEM for 100 nm, 200 nm, 300 nm PS beads embedded in 0.12%, 0.23%, 0.47%, 0.94%, 1.88%, 3.75%, 7.5% of poloxamer 407.*

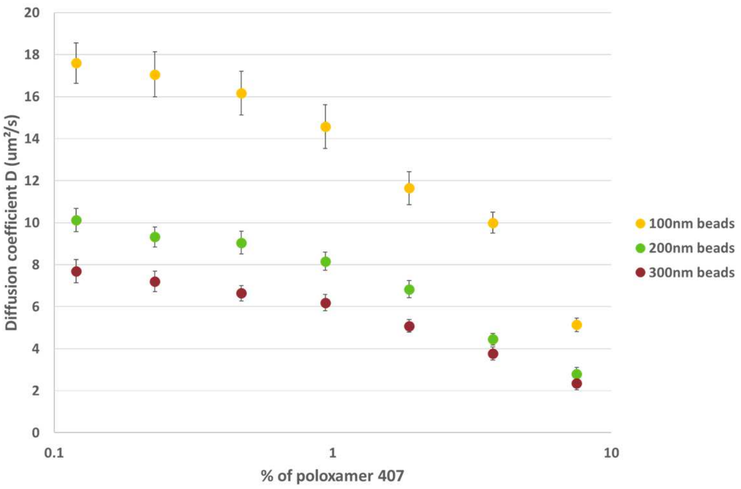


*Supplementary Figure 6: Diffusion coefficient of 100 nm, 200 nm and 300 nm PS beads as a function of the percentage of poloxamer 407.*
